# Variable jackpot individuals provide most alleles for repeated, rapid adaptation to freshwater by anadromous Threespine Stickleback

**DOI:** 10.64898/2026.05.29.728866

**Authors:** Alexander Kwakye, Mark K. Sanda, Krista Oke, David C. Heins, Michael A. Bell, Kerry Reid, Krishna R. Veeramah

## Abstract

Experimental introductions of anadromous stickleback into freshwater habitats lacking this species allow analysis of the process of adaptation to freshwater forward-in-time. We examined the population genomic dynamics during early stages of adaptation in three replicate lakes that were experimentally founded, each using ∼3000 anadromous ancestors. We replicated earlier results that rare individuals carrying large haploblocks of freshwater-adaptive alleles (jackpot carriers) provide most of the allelic variation for adaptation of anadromous Threespine Stickleback to freshwater within only a few generations in each lake. There were population bottlenecks two to three generations after founding in each lake, after which jackpot carriers dramatically increased in frequency and came to dominate the populations. Individuals lacking large adaptive haploblocks experienced low fitness in their new freshwater environments, consistent with our previous report based on a single lake population. Despite similarities of the demographic responses to selection, the alleles that were most common among jackpot carriers were different in each population, suggesting that each lake population likely adapted to conditions in freshwater environments through different genes. These results provide direct evidence for the genomic mechanisms underlying the rapid adaptation of anadromous sticklebacks to freshwater environments, a process that can occur within just a few generations.

## Introduction

There are numerous examples of rapid adaptation resulting from selection on standing genetic variation (SGV) (Cresko et al. 2004; Colosimo et al. 2005; Franks et al. 2007; Barrett and Schluter 2008; Pritchard et al. 2010; Hermisson and Pennings 2017; Schrider and Kern 2017; Roberts Kingman et al. 2021). In particular, *Gasterosteus aculeatus* (Threespine Stickleback) has emerged as an important model to study the dynamics of adaptation from SGV in vertebrate species (Reid et al. 2021). The ancestral oceanic (marine and anadromous) ecotype has repeatedly colonized freshwater for at least the past 10 million years (Bell and Foster 1994; Bell 2009). Adaptation of the colonizing populations to new freshwater environments is characterized by rapid and repeated phenotypic changes occurring within a few generations (Heuts 1947; Munzing 1963; Bell and Richkind 1981; Klepaker 1993; Bell 2001, 2013; Bell et al. 2004; Aguirre et al. 2022).

Previous studies have identified more than 300 loci with alleles that have been continually reused during divergence of freshwater populations from oceanic ecotypes across the range of Threespine Stickleback (Cresko et al. 2004; Colosimo et al. 2004; Jones et al. 2012; Marchinko et al. 2014; Roberts Kingman et al. 2021). These loci form highly linked haplotype structures within which there are multiple beneficial alleles (Roberts Kingman et al. 2021). These loci tend to be interspersed by local recombination hotspots that potentially facilitate their reassembly into larger, concentrated haploblocks that maximize fitness during freshwater adaptation which are observed in long-established freshwater populations (Schluter and Conte 2009; Yeaman 2022). However, given the rapid speed with which freshwater adaptation has been observed to take place (Bell et al. 2004; Roberts Kingman et al. 2021; Aguirre et al. 2022), until recently it remained unclear how recombination was able to reassemble such large haploblocks in the first few generations of colonization. Most oceanic stickleback only appear to possess a few freshwater-adaptive alleles on an individual by individual basis (Jones et al. 2012; Roberts Kingman et al. 2021). At the same time, recombination rate observed along much of the chromosome containing these adaptive loci are very low, with previous work showing that as many as half of all chromosomes in males, as well as a third of chromosomes in females, are likely to be inherited without any recombination event each meiosis (Venu et al. 2024).

By analyzing approximately 350 whole genomes from the earliest founding generations of the recently introduced Scout Lake population in Alaska, we provided evidence that adaptation of anadromous stickleback to the new freshwater environment can be mediated predominantly by a few individuals with large haploblocks of freshwater-adaptive alleles, referred to as “jackpot carriers” (Kwakye et al. 2026b). These jackpot carriers appear to be present in the oceanic population at low frequencies (<0.1%) and possess at least 5% of the full complement of freshwater alleles, likely reflecting a mixture of circulating first to third-generation anadromous-freshwater hybrids. The majority of individuals carrying few freshwater-adaptive alleles exhibited low fitness when restricted to freshwater environments, resulting in a population crash within a few years after colonization. Consequently, non-jackpot individuals contributed few descendants to subsequent generations, except through interbreeding with jackpot carriers. In contrast, positive selection favored the low frequency jackpot carriers, allowing the population to recover from this crash and eventually grow and establish, such that most individuals in the lake after six years were connected within a close kinship network.

In this study, we used whole genome resequencing of samples collected from two additional founded freshwater lake populations (Bell et al. 2016; Aguirre et al. 2022; Kwakye et al. 2026b), to investigate the extent to which the dynamics observed in Scout Lake were part of a more general process of freshwater adaptation for Threespine Stickleback.

## Results

### Resequencing of whole genomes from newly founded populations

We generated new low coverage TN5-based whole genome resequencing data (n=131) from Warfle Lake (three, four and five years after founding, see supplementary note for details on founding of stickleback population in this lake) and Cheney Lake (six years after founding) (Fig. 1). We refer to the Warfle samples as WL2022 (n=16, mean coverage=0.58X), WL2023 (n=37, mean coverage=1.43X) and WL2024 (n=54, mean coverage=1.82X) to indicate samples collected three, four and five years after founding, respectively, while the Cheney Lake sample is labelled CH2015 (n=24, mean coverage=2.6X). We also analyzed these new sequences alongside previously published genome sequences. The previously published genomes included sequence data collected two, three, four, six and nine years after the Scout Lake population was founded and refer to them as SC2013, SC2014, SC2015, SC2017 and SC2020, respectively; and a sample from 11 years after founding of Cheney Lake (CH2020) (Kwakye et al. 2026b). Samples from Rabbit Slough collected in 2009 (RS2009) (Roberts Kingman et al. 2021) and 2019 (RS2019) are used to represent genomic variation in the founding populations (Kwakye et al. 2026b). In total, we analyzed 623 genomes consisting of 492 previously published genomes (Fig. 1, Table 1).

**Fig. 1.**
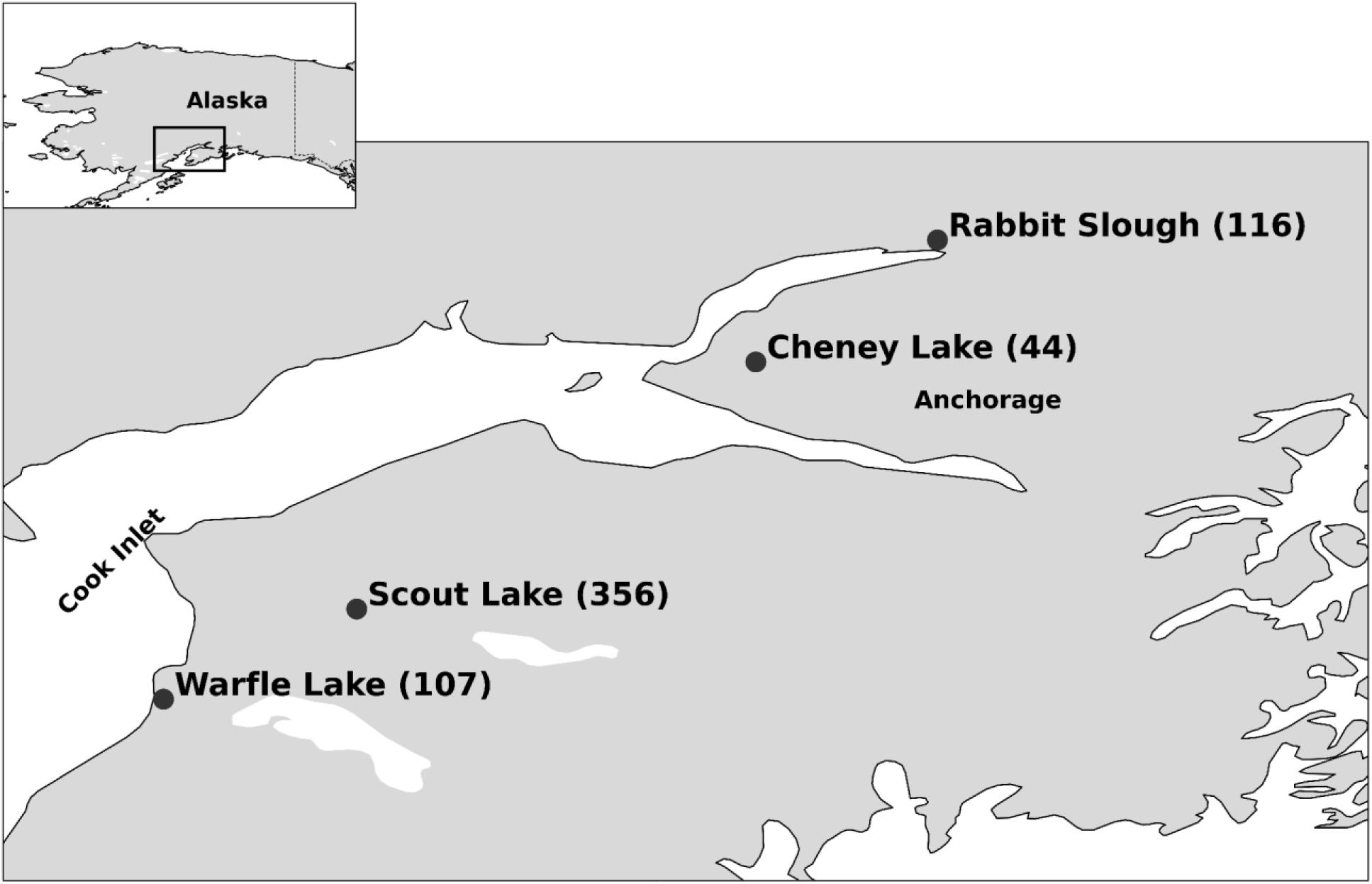
Locations of newly founded freshwater Threespine Stickleback populations in Cheney, Scout and Warfle lakes and the source of anadromous stickleback (Rabbit Slough) used to found them. Numbers in parentheses indicate the number of genomes from each location.

**Table 1:** Sample sizes and mean sequencing coverage of selected samples from Rabbit Slough (RS), the ancestral population, and from descendant populations in Scout (SC), Cheney (CH) and Warfle (WL) lakes. Lake acronyms in the Samples column are followed by the year in which the sample was made.

| Samples | Sample size | Mean coverage |
| --- | --- | --- |
| RS2009 | 20 | 30X |
| RS2019 | 96 | 1.84X |
| SC2013 | 96 | 1.31X |
| SC2014 | 48 | 0.43X |
| SC2015 | 96 | 0.78X |
| SC2017 | 96 | 1.01X |
| SC2020 | 20 | 25.7X |
| CH2015 | 24 | 2.6X |
| CH2020 | 20 | 30X |
| WL2022 | 16 | 0.58X |
| WL2023 | 37 | 1.43X |
| WL2024 | 54 | 1.82X |
| Total | 623 |  |

### Determining putative cohorts in the initial years after founding

The short growing season available in Cook Inlet lakes allows only a short, early breeding season because fish born late in the growing season are less likely to accumulate enough resources to overwinter (Cargnelli and Gross 1996). In addition, fish may also breed earlier in years when the ice melts early as a result of climate change, which leads to a longer breeding season (Hovel et al. 2017). A previous study of Scout Lake stickleback indicated that the stocked adults likely did not survive the winter after they were released into the lake (Kurz et al. 2016). In addition, no individuals with an anadromous phenotype have been observed in samples from the year after release of anadromous stickleback in any of the three lakes. Sticklebacks that were caught in the second year were larger and less common because they had a second year of normal attrition, and the second generation in the lake (F2) had not bred in large numbers yet.

The one-year-old F1 stickleback in Scout Lake probably did not reproduce until they reached two years old (Kurz et al. 2016; Kwakye et al. 2026b). Therefore, the samples collected three years after founding in Scout Lake (SC2014) were probably one-year old F2 (Fig. 2A), which likely did not become sexually mature until they were two years old in 2015. Based on analyses of standard lengths of samples collected in the first three years after the Scout Lake population was founded, we inferred the likely predominant generation in each sampled year for all three lakes (Kurz et al. 2016) (Fig. 2A-C).

**Fig. 2:**
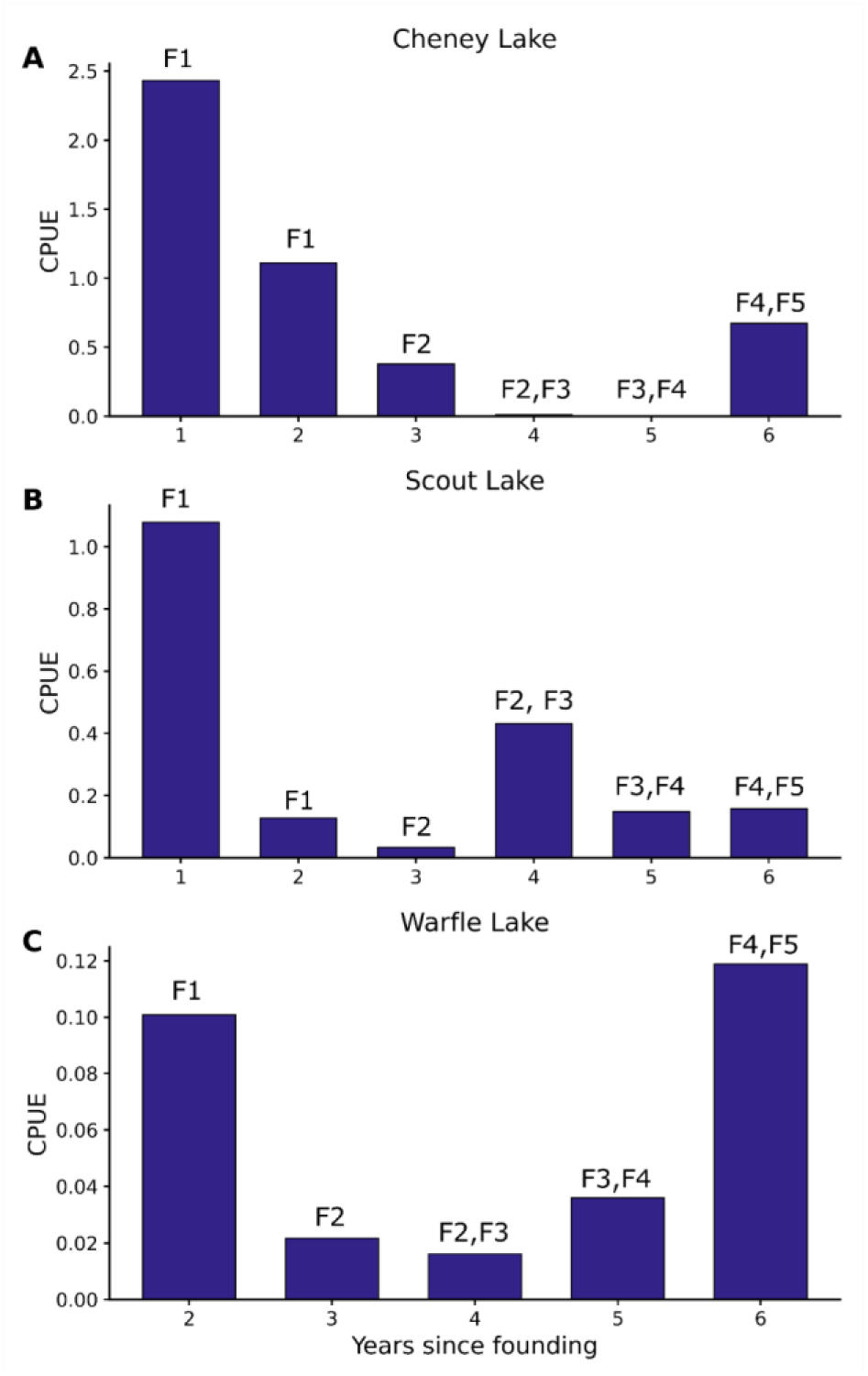
Mean catch per unit effort (CPUE) across the three study lakes and inferred predominant generations. CPUE estimates are shown for samples collected from (A) Cheney Lake (founded in 2009), (B) Scout Lake (founded in 2011), and (C) Warfle Lake (founded in 2019). In all panels, labels above the bars (e.g., F1, F5) indicate the inferred predominant generation represented in each sample. Bars represent mean CPUE values calculated from replicate samples collected at each time point. For Cheney Lake, the number of sampling events was *n* = 8, 9, 15, 4, 1, 5 for 1–6 years since founding, respectively. For Scout Lake, the number of sampling events was *n* = 4, 13, 4, 5, 1, 1 for 1–6 years since founding, respectively. For Warfle Lake, the number of sampling events was *n* = 1 for all sampled time points. The number of sampling events indicates the number of times that sampling was attempted during the breeding season.

The Cheney Lake population was founded in 2009, and the samples collected in 2010 were F1 (i.e., first generation born in the lake), with no evidence of breeding as one year olds. As in Scout Lake, Cheney Lake F1s appeared to breed in 2011 at two years old. In 2012, three years after the population was founded, most of the fish caught in Cheney Lake were likely one-year-old F2 fish (Fig. 2B). The F2 and F3 were mostly present in the samples collected in 2013 and 2014, four and five years after founding, respectively, while the 2015 sample (CH2015) likely consisted of F3 and F4 individuals (Fig. 2B).

Warfle Lake was founded in 2019, with the first summer collection sampled in 2021. The sample collected in the 2021 likely consisted mostly of two-year-old F1 stickleback, while the 2022 sample (WL2022) were therefore one-year-old F2 fish. The 2023 (WL2023) and 2024 (WL2024) probably consisted mostly of F2 and F3 fish, while the sample collected in 2025 was likely dominated by F3 and F4 (Fig. 2C).

### Demographic bottleneck during early generations

To determine the temporal dynamics of population density in the three newly founded lakes, we estimated the catch per unit effort (CPUE) for each sampling year (Fig. 2A-C) (Bell et al. 2016). CPUEs were estimated from samples collected during the breeding season. In Scout Lake, there was a population density decline between the one-year old F1 and two-year old F1 (mean CPUE in 2012=1.08 fish per trap-hour, 2011=0.127 fish per trap-hour, Fig. 2B). The population density declined further in 2014, when putative one-year old F2 were sampled (mean CPUE= 0.033). The following year, in 2015, the population density increased (mean CPUE=0.43 fish per trap-hour) (Bell et al. 2016; Kwakye et al. 2026b).

In both Cheney and Warfle, as in Scout Lake, population density decreased dramatically after the F1 generation (Fig. 2) and recovered in subsequent generations. In Scout and Warfle lakes, the populations recovered in the fourth year. However, the population density remained low in years four and five in Cheney Lake, and only recovered in the sixth year after founding. These results point to consistent demographic patterns in the first few generations after the founding of all three freshwater populations, although the specific year of the population crash after founding differed from lake to lake.

### Genotypes of freshwater-adaptive loci

We characterized the temporal dynamics of freshwater alleles during freshwater adaptation in the three lakes. The genotypes at multi-SNP haplotypes of freshwater-adaptive alleles previously identified by Roberts Kingman et al. (2021) were estimated using an approximate genotype likelihood approach developed in Kwakye et al. (2026b). There were 341 genome-wide multi-SNP haplotypes, each made up of between three to 3,658 SNPs, with a median size of 27.3 kb. Out of the 341 loci, we filtered out any locus where an individual had a missing genotype, resulting in 280 loci with no missing data. All downstream analyses were based on these 280 loci. We determined the genotype at each locus as homozygous oceanic, heterozygous, or homozygous freshwater and defined individuals with >5% freshwater–adaptive alleles as jackpot carriers Kwakye et al. (2026b).

No jackpot carriers were observed in the ancestral Rabbit Slough population in either 2009 or 2019 samples (Kwakye et al. 2026b). The number of jackpot carriers increased dramatically in the Scout Lake population from 1% in SC2013 (two years after founding) to about 50% in SC2014 (three years after founding, mostly F2). By the sixth year after founding, all the sampled individuals were descendants of jackpot carriers. The number of homozygous freshwater loci also increased over time, which likely reflected the mating between the predominantly heterozygous descendants of jackpot carriers (Fig. 3E).

**Fig. 3:**
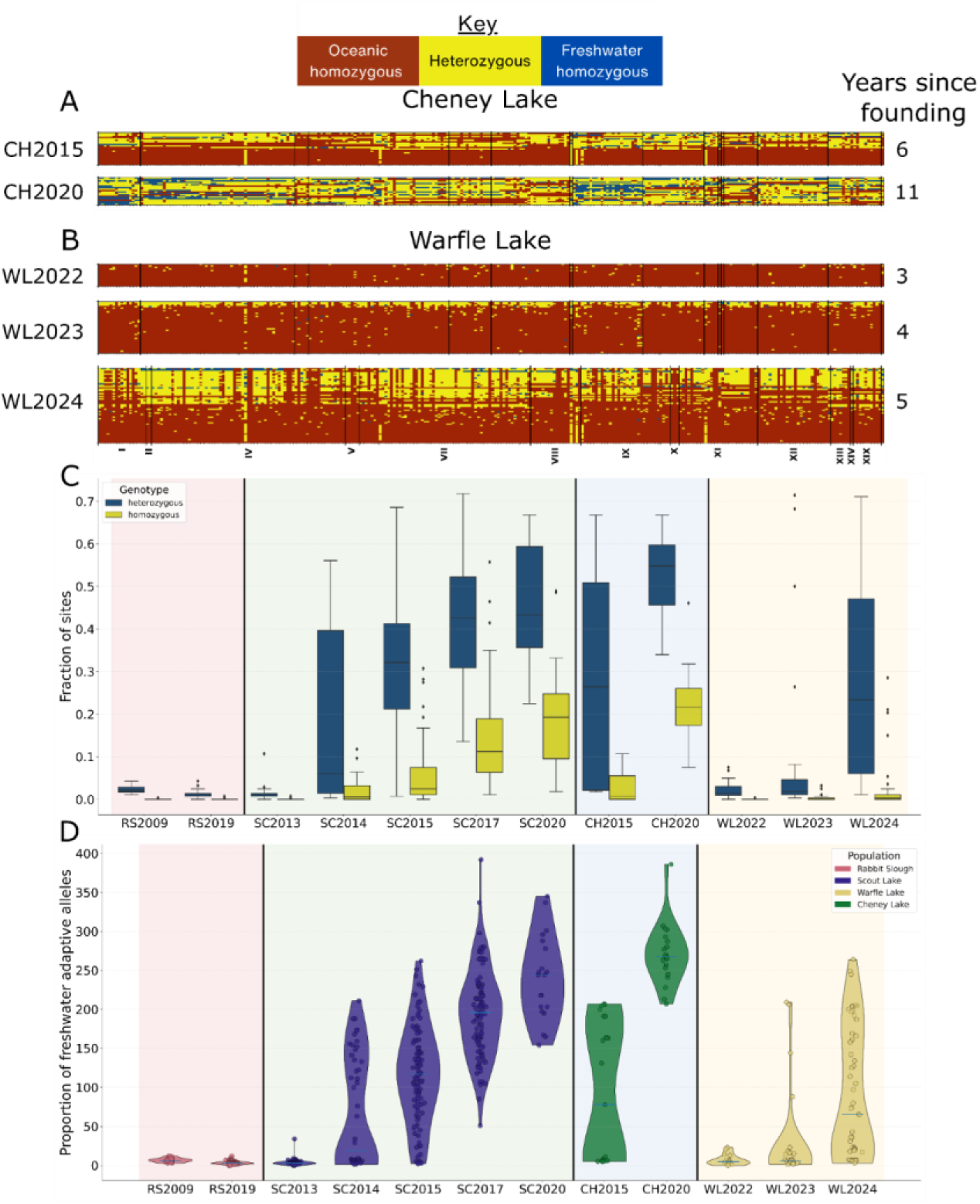
Genotypes at freshwater adaptive loci. **A)** Genotypes of individuals from Cheney Lake samples 6 and 11 years after its founding in 2009. **B)** Genotypes of individuals from Warfle Lake collected three, four and five years after its founding in 2019. Comparable plots for Scout Lake samples are in Figure 1, Kwakye et al. (2026b). **C)** The fraction of loci that are heterozygous and homozygous for the freshwater allele in each timepoint. **D)** The proportion of freshwater alleles at all 280 freshwater-adaptive loci across all timepoints in Rabbit Slough and the descendant lake populations. In panels A and B, each row is an individual and each column is one of the 280 loci and individuals are ordered from those that carry the most freshwater alleles to least in each plot. The sample size for CH2015=24, CH2020=20, WL2022=16, WL2023=37, WL2024=54. Each locus is defined by a multi-SNP region consisting of three to 3658 SNPs. Panels C and D include samples from the ancestral, anadromous, Rabbit Slough population (RS2009 and RS2019) Scout Lake (SC2013, SC2014, SC2015, SC2017, SC2020; Kwakye et al. (2026b)

There were 12 descendants of jackpot carriers among 23 individuals from CH2015, while all the 20 individuals in CH2020 were descendants of jackpot carriers. There were no jackpot carriers in WL2022 (n=16), four in WL2023 (n=37) and 32 in WL2024 (n=54). The years that descendants of jackpot carriers (putative F2 and F3 generations) first appeared in our samples were one or two generations after the start of the bottleneck in all three populations. The large haploblocks present in these initial descendants of jackpot carriers were also predominantly heterozygous, with freshwater homozygotes appearing only in a later generation in Scout and Cheney lakes. Warfle samples contain limited homozygotes because fewer generations have been sampled since founding (Fig. 3C). These results indicate that the descendants of jackpot carriers are likely to have been selectively favored in the new freshwater environments. The increased fitness of jackpot carriers in the freshwater environments was likely because they possessed many freshwater-adaptive alleles that permitted them to breed and/or survive. In contrast, descendants of the non-jackpot individuals possessed few freshwater-adaptive alleles and were less fit, reducing the relative frequency of non-freshwater-adaptive alleles and initiating the demographic bottlenecks indicated by CPUE results.

### Biological relatedness in Cheney and Warfle Lakes

Kinship analysis of stickleback samples from Scout Lake showed that the samples collected after 2014 consisted of many closely related individuals that were descended from the jackpot carriers (Kwakye et al. 2026b). By the sixth generation (i.e., F6) in Scout, every individual sampled was a descendant of the set of initial rare jackpot carriers present in the founders, indicating that their high reproductive success drove both the population’s demographic recovery and a dramatic increase in freshwater-adaptive alleles derived from the jackpot carriers. Given that we observed similar demographic trends and freshwater-adaptive allele frequency increases in the Cheney and Warfle Lake populations, we investigated biological relatedness among individuals in these populations using READv2, which estimates biological relatedness up to third-degree relatives in low coverage sequence data (Alaçamlı et al. 2024).

In Cheney Lake, there were two pairs of second-degree relatives and three pairs of third-degree relatives in the CH2015 sample, while there were two pairs of second-degree and five pairs of third-degree relatives in the CH2020 sample. There were also seven third-degree relatives in which one member of the pair was in CH2015 and the other in CH2020. All but two pairs of relatives were between jackpot carriers (Fig. 4A). Considering the small size of CH2015, these results are largely consistent with the general kinship pattern observed in Scout, though interestingly, we observed no related pairs in the Scout 2020 sample analyzed by Kwakye et al. (2026b), despite having the same sample size as CH2020. Considering that it was founded two years earlier than Scout, this suggests a somewhat smaller effective population size in Cheney.

**Fig. 4:**
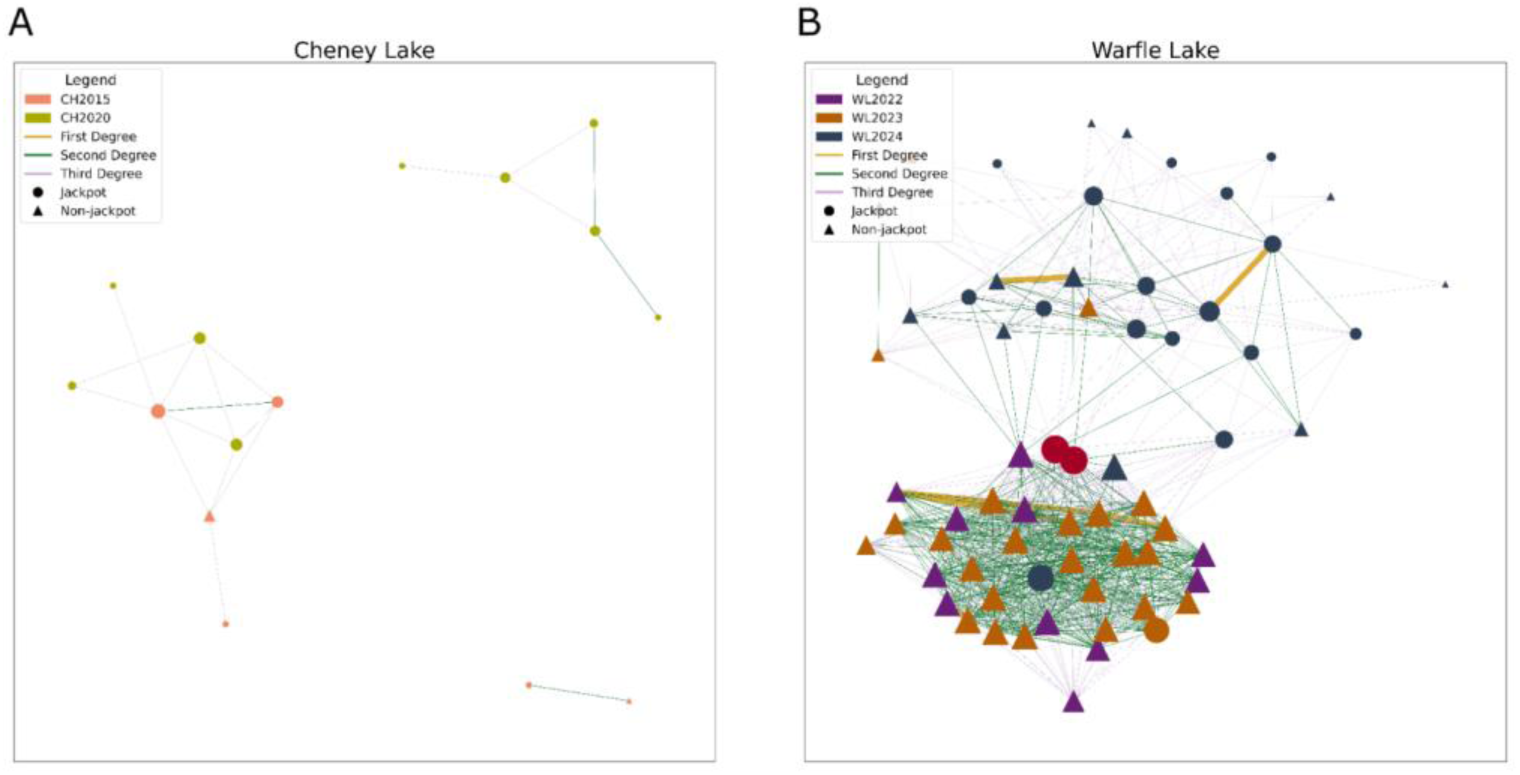
Biological relatedness between specimens from Cheney and Scout Lakes. **A)** Samples from Cheney Lake are CH2015 and CH2020, and those from **B)** Warfle Lake are WL2022, WL2023 and WL2024. Each node represents one individual and the edges represent the number of individuals to which its node is connected by first- to third-degree relationships. Larger nodes have more connections.

In Warfle Lake we observed six first-degree, 488 second-degree and 287 third-degree related pairs (Fig.4B). Three out of the six first-degree relatives were siblings with one of the pair from WL2022 and the other from WL2023. There were two pairs of first-degree relatives that were assigned as possible parent-offspring pairs from WL2024, demonstrating that WL2024 consisted potentially of more than one generation as inferred above. There was however insufficient support to distinguish the second pair of first-degree relatives as either parent-offspring or full siblings. All of these pairs (first-, second- and third-degree) formed a single large network consisting of individuals within and across the three timepoints (Fig.4B). We observed that one out of the four, and 18 out of the 32 jackpot carriers in WL2023 and WL2024, respectively, were part of this large network of relatedness. The network was divided into two subnetworks, one consisting of individuals from WL2022 and WL2023, and the other consisting of individuals from WL2024. Notably two jackpot carriers from WL2024 (colored red in Fig. 4B) had the most connections in the network (40 out of 65 nodes in the network) (Fig.4B) and connected these two subnetworks. We also observed that one of the subnetworks consisted predominantly of non-jackpot individuals from WL2022 and WL2023, i.e. prior to the demographic bottleneck. It should be noted that we did not observe such a large network of related individuals prior to the equivalent bottleneck in Scout Lake.

### Population specific haploblocks

To assess if certain freshwater alleles were repeatedly enriched in jackpot carriers during the earliest stages of the adaptive process, we determined freshwater allele frequencies in jackpot carriers in each population. In Scout Lake, the locus with the greatest frequency of freshwater-adaptive alleles was a 22Kb region on chromosome IV (chrIV:14326747–14349257) (Fig. 5, Supplementary data), which was present in 170 out of 225 jackpot carriers. Notably, all 29 loci with freshwater adaptive alleles in the 90th percentile were located on chromosome IV (Table 2).

**Fig. 5:**
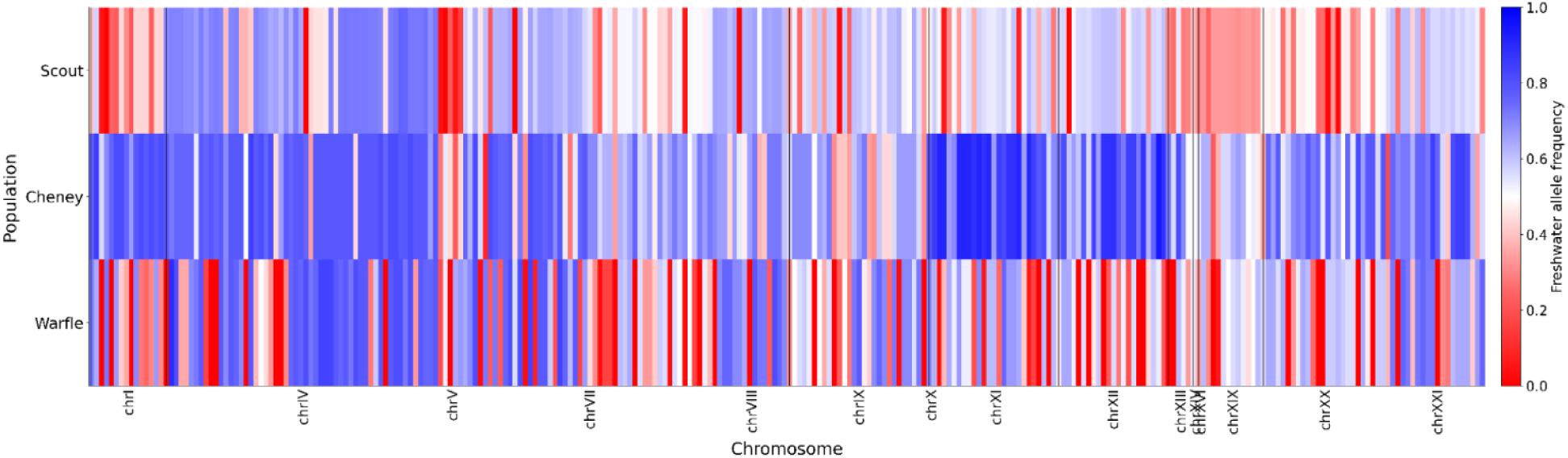
Different loci are enriched with freshwater adaptive-alleles in different populations. Freshwater adaptive allele frequencies across all jackpot carriers in the three lakes across all 280 loci.

**Table 2:** Number of loci with alleles in the 90th percentile of most frequent freshwater-adaptive alleles for each population. For each population, we group the alleles by chromosomes.

| Chromosome | Scout | Cheney | Warfle |
| --- | --- | --- | --- |
| chrI | 0 | 1 | 0 |
| chrIV | 29 | 1 | 23 |
| chrVII | 0 | 0 | 11 |
| chrXI | 0 | 16 | 0 |
| chrXII | 0 | 9 | 0 |
| chrXXI | 0 | 3 | 0 |

However, in Cheney Lake, the most frequent freshwater adaptive allele was found in a 23Kb region found on chromosome XII (chrXII:4384523–4408327), which was observed in 30 out of the 32 jackpot carriers (Fig. 5, Supplementary data). Unlike in Scout Lake, the majority of the loci in the 90th percentile with the highest freshwater-adaptive allele frequencies were found on either chromosome XI (n=16) or XII (n=9) (Table 2). Of these loci, only one was found on chromosome IV and none on chromosome VII. None of the loci with freshwater-adaptive alleles in the 90th percentile were shared between Cheney and Scout lakes (Supplementary data).

In Warfle Lake, a 16Kb region on chromosome IV (chrIV:24441665–24457941) possessed the most frequent freshwater-adaptive alleles, which occurred in 32 of 36 jackpot carriers (Table 2). Out of 34 loci with freshwater-adaptive alleles in the 90th percentile, 23 (68%) were located on chromosome IV, while the remaining ones were found on chromosome VII (Table 2). There was one locus (chrIV:27357302–27374317) shared between Warfle and Cheney lakes, and 17 loci (all on chromosome IV) shared between Warfle and Scout Lakes (Supplementary data).

The correlation of freshwater allele frequencies among the populations was generally low (Table 3). For example, the strongest correlation between allele frequencies in jackpot carriers only in Scout and Cheney lakes was *r*=0.27 (p-value = 3.33e-06; Table 3, lower triangle). The correlation between allele frequencies estimated from jackpot carriers in the Cheney and Warfle populations was approximately half of the other pairwise correlation coefficients (*r*=0.13, p-value = 0.027, Table 3). The adaptive region that contains the *EDA* gene on Chromosome IV had freshwater alleles in 70% (92th percentile, 158 out of 225), 81% (89th percentile, 26 out of 32) and 78% (90th percentile, 28 out of 36) of jackpot carriers in Scout, Cheney and Warfle Lakes, respectively.

**Table 3:** Pairwise correlation matrix using freshwater adaptive allele frequencies across all 280 loci. The top triangle was estimated using allele frequency from all individuals and the lower triangle was from allele frequencies from only jackpot carriers.

|  | Warfle | Scout | Cheney |
| --- | --- | --- | --- |
| Warfle |  | 0.21 | 0.25 |
| Scout | 0.26 |  | 0.28 |
| Cheney | 0.13 | 0.27 |  |

We next identified genes overlapping the adaptive loci with most frequent freshwater alleles in each lake. In Scout Lake, the locus with the most frequent freshwater allele was a 22Kb region located on chromosome IV, which overlapped two genes (*foxi3b* and *wnt8a*). A 23Kb region found on chromosome XII in Cheney Lake overlapped ENSGACG00000003535 and *ovgp1* while the 16Kb region on chromosome IV in Warfle Lake did not overlap any gene.

Previous studies have suggested that *cis*-regulatory changes that alter gene expression may likely be the dominant genetic mechanism underlying freshwater adaptation (Jones et al. 2012; Mack et al. 2023; Kwakye et al. 2026a). To capture putative *cis*-regulatory regions that overlapped the loci with the most frequent-adaptive alleles, we searched for genes within 10Kbp up- and down-stream of these regions. In Scout Lake, the 22Kb region located on chromosome IV overlapped the *cis*-regulatory regions of *afap1l1a* and *gabrp;* the locus with the most frequent-adaptive allele in Cheney Lake overlapped ENSGACG00000003505, ENSGACG00000012566, and ENSGACG00000012562; and the locus with the most frequent freshwater-adaptive allele in Warfle Lake overlapped *pfkfb3*. Gene ontology analyses did not show any enriched pathways among the genes overlapping the loci with freshwater-adaptive alleles in the 90th percentile across the three lakes.

## Discussion

### Jackpot carriers mediate rapid freshwater adaptation

In this study, we show that adaptation to three distinct freshwater environments by anadromous sticklebacks was mediated by strong positive selection favoring a few individuals with large haploblocks of freshwater-adaptive alleles (i.e. jackpot carriers, (Bassham et al. 2018)) present in the founders. In all three populations, there were demographic bottlenecks in the first few years that resulted from reduced fitness of a majority of the individuals possessing only a few freshwater-adaptive alleles. Interestingly, we observed very different kinship dynamics amongst each of the three populations, which may reflect differences in fitness of founding jackpots carriers. Indeed, while there were similar overall demographic trends across all three lakes, the jackpot carriers in each lake population had different profiles of freshwater-adaptive alleles, suggesting that each lake population adapted to conditions in freshwater environments through different combinations of genes.

The year after the introductions in each lake, the F1 progeny of the founders were abundant; but catch per unit effort declined sharply in the F1 cohort’s second year. This decline likely reflected normal attrition during the first year of life, which causes collections from established lake populations to contain many more one-year-old sticklebacks than two-year olds. There is no reason to expect this mortality to have been selective because anadromous sticklebacks spend their first two months of life in freshwater. Remaining in freshwater may, however, deprive descendants of anadromous sticklebacks of nutrients that they would have acquired in the ocean (Ishikawa et al. 2019), interfere with their reproduction, and decrease population density that we observed during the first year of the F2 generation in the lakes (Bell et al. 2016). The F2 generation was the same generation in which the frequency of freshwater-adaptive alleles first increased (Roberts Kingman et al. 2021) and haplotypes with numerous freshwater-adaptive alleles first appeared (Kwakye et al. 2026b; this study). Catch per unit effort tended to increase during subsequent generations and frequencies of many freshwater-adaptive alleles started to increase progressively among generations (Roberts Kingman 2021; this study). It appears that population growth after the F2 generation in all three lakes resulted from increasing frequencies of individuals with large haploblocks containing numerous freshwater-adaptive alleles (Kwakye et al. 2026b; this study).

We observed two lake-specific phenomena: population specific freshwater-adaptive allele-frequency changes (Fig. 5) and kinship dynamics. These lake-specific observations may reflect differences in fitness of the descendants of jackpot carriers, absence of some alleles among the founders in different lakes, different genetic backgrounds that changed the fitness effects of freshwater-adaptive alleles through epistasis or ecological differences among the three lakes. We note that Warfle and Scout Lakes are proximal to each other and may share similar ecological variables, which possibly resulted in an increased number of loci with freshwater-adaptive alleles in the 90th percentile shared between them compared to Cheney Lake. These results also indicate that there are likely to be many combinations of freshwater-adaptive alleles that can be the basis for adaptation of anadromous stickleback to freshwater, and it is the proportion of freshwater alleles present in the jackpot carriers sampled to found the population rather than the specific configuration that is important. There is evidence that replicate populations could adapt to similar environmental conditions through different genes (Hoekstra and Nachman 2003; Conte et al. 2015; Bolnick et al. 2018)

In Scout Lake, the genes forkhead box I3b *(foxi3b*) and *Wnt* family member 8A (*wnt8a*) overlapped the loci with the most frequent freshwater-adaptive allele. The gene *foxi3b* is a transcription factor that is involved in ionocyte differentiation (Janicke et al. 2010; Cruz et al. 2013; Hwang and Chou 2013; Chen et al. 2021), while *wnt8a* is involved in dorsoventral and anteroposterior patterning during development (Baker et al. 2010). *Wnt* signalling is suggested to be involved in armor plate development in sticklebacks by acting upstream of *EDA (O’Brown et al. 2015)*. ENSGACG00000003535, which was one of two genes overlapping a region with greatest frequency increase in the Cheney Lake population, is predicted to be a chitinase, which may be involved in innate immunity during development in zebrafish (Teng et al. 2014). Oviductal glycoprotein 1 (*ovgp1),* the other gene that increased rapidly in frequency in the Cheney Lake population is a major component of the oviductal fluid, involved in gamete maturation and fertilization (Buhi 2002). This gene also has chitinase-like domains (Choudhary et al. 2019). *Pfkfb3*, which is located within 10Kb of the loci with the most frequent freshwater-adaptive allele in the Warfle Lake population, is involved in glycolysis and has been found to be differentially expressed in the brain of a long-established freshwater stickleback population (Kwakye et al. 2026a). Thus, despite high frequencies of different freshwater-adaptive alleles in each population, it appears that the majority of the genes associated with them are involved in development. Development associated genes such as *EDA (O’Brown et al. 2015)* may be critical for freshwater adaptation, and suggest that the strength of natural selection early in life may outweigh possible late life counteracting selective pressures (Barrett et al. 2008).

### Founder effects during establishment of new populations

The adaptation of anadromous sticklebacks to the three replicate freshwater environments allow insights into the evolutionary processes underlying successful establishment in novel environments. For example, the evolutionary consequences of founding population size are central to ecological theory, conservation practices (Willis and Willis 2010), and theories of speciation such as the founder effect–genetic revolution model (Mayr 1954, 1999), flush-crash model (Carson 1975) and genetic transilience model (Templeton 1980). In all three lake populations, a small subset of the founders (jackpot carriers) contributed most of the freshwater-adaptive alleles as a result of the reduced fitness of the majority of individuals that bottlenecked the populations, but the rapid spread of alleles was due to their mating with non-jackpot individuals. In his discussions of phenotypic divergence and speciation, Ernst Mayr emphasized the significance of demographic bottlenecks that result in a few individuals as the foundation for new populations. This reduction in population size then leads to loss of heterozygosity through genetic drift (founder effect), essentially changing the genetic background (so-called genetic revolution) (Mayr 1954, 1999; Barton and Charlesworth 1984). The adaptation to the three replicate freshwater environments appears to have involved founder effect, although the reduction in population size was likely driven by both non-selective (genetic drift) and selective processes (selection on jackpot carriers).

Founder effects and genetic revolutions also underpin other speciation theories like Carson’s flush-crash model, which divided the genome into an open system and a closed system (Carson 1975). The open system consists of recombining genomic regions with limited epistasis, which underlie quantitative traits as well as facilitate gene flow, whereas the closed system comprises tightly linked, coadapted genes with strong epistasis that resist gene flow and preserve adaptive haplotypes. The haploblock of freshwater-adaptive alleles that were present in jackpot carriers constitute such a closed genetic system with accumulated beneficial alleles in low recombination regions (Roberts Kingman et al. 2021; Venu et al. 2024). According to the flush-crash model, reorganization of the closed genetic system occurs through a series of stochastic genetic events and leads to speciation. However, such reorganizations are only possible during a phase of population growth (flush phase, (Carson 1968, 1971)) from relaxed selection pressure and an increased survival of recombinants. Although the instances of adaptation of anadromous stickleback to the three freshwater environments involved drastic changes in the gene pool through bottlenecks (crash phase) followed by a population growth (flush phase), these changes were largely observed in the open genetic system. In addition, the flush phase, according to the theory, should be accompanied by limited selection pressure. Thus, the crash-flush phases and the accompanying reorganization of the gene pool during freshwater adaptation in three replicate lake populations may not fit the predictions of the flush-crash theory.

With regards to founding population size, there is also evidence from the literature that rapid adaptation has proceeded from a few founders in other systems. In *Daphnia*, as few as five founders have been suggested to establish standing genetic variation required for rapid evolution in natural populations of the waterflea (Chaturvedi et al. 2021). In the saltmarsh beetle, *Pogonus chalceus,* there is evidence that the tidal ecotype could evolve from approximately five to fifteen long-winged seasonal ecotypes (Van Belleghem et al. 2018). An accidental introduction of approximately 72 pink salmon (*Oncorhynchus gorbuscha*) was enough to establish this species in the Great Lakes (Sparks et al. 2024) (see also (Langan et al. 2026)). These cases provide support for successful establishment of new populations to novel environments from only a few founders through standing genetic variation.

### Maintenance of standing genetic variation required for rapid freshwater adaptation

The freshwater-adaptive alleles required for the successful establishment of stickleback populations to the new lakes were likely maintained in the ancestral range through gene flow – selection balance, as proposed by the transporter hypothesis (Schluter and Conte 2009).

Anadromous and freshwater stickleback often breed simultaneously in sympatry and produce hybrids (Hagen 1967; Karve et al. 2008), so hybridization is a plausible source of freshwater-adaptive alleles in anadromous stickleback. Aside from gene flow–selection balance, standing genetic variation could be maintained in the ancestral range either through mutation–drift balance, selective neutrality (Haenel et al. 2022), some form of balancing selection (Stern and Lee 2020; Ruzicka et al. 2026) or a combination of these. Beneficial alleles involved in adaptation to multiple acidic freshwater habitats in the North Uist segregate neutrally in the marine environment (Haenel et al. 2022). Stern and Lee (2020) showed that in the saltmarsh copepod, *Eurytemora affinis,* multiple instances of adaptation to freshwater habitats involved loci with balanced variants as a result of fluctuating selection in the native populations. Other forms of balancing selection, such as dominance reversal (Grieshop and Arnqvist 2018), antagonistic pleiotropy (Hedrick 1999) or multiple forms of these (Paris et al. 2025) could also maintain standing genetic variation in the ancestral environment. Future studies could explore the extent to which gene flow–selection balance maintains freshwater-adaptive alleles in the ancestral marine environments, relative to other mechanisms.

## Conclusion

This study confirms previous reports that extremely rapid adaptation of anadromous Threespine Stickleback to freshwater depends on jackpot carriers, individuals that possess dozens of freshwater adaptive alleles probably because they are hybrids or descendants of a recent hybridization event between an anadromous and a freshwater parent. Most of the alleles for adaptation to freshwater come from a small number of jackpot carriers with high fitness in freshwater. However, the frequencies of different freshwater-adptive alleles increased rapidly among the jackpot carriers in the three lakes. These results show that the outcomes of adaptation are influenced both by the genetic composition of the founders and the local ecological environments facilitating rapid adaptation to freshwater.

## Methods

### Founding of the lake populations

The methods to found the populations in Cheney Lake in 2009 and Scout Lake in 2011 were described by (Bell et al. 2016; Aguirre et al. 2022; Kwakye et al. 2026b). All fish used in this study were collected according to an approved protocol from the Institutional Animal Care and Use Committee (IACUC 1446584) at Stony Brook University. We describe the general stocking methods that established the populations here briefly.

Anadromous Threespine Stickleback were captured in Rabbit Slough either at a culvert under Glenn Highway (61.536N, 149.253W) or within about 100 m of the junction of Rabbit Slough and Spring Creek (61.534N, 149.265W), and about 1 km downstream from the culvert. The stickleback were transported live in coolers of aerated ambient water, held overnight in pools at the University of Alaska Anchorage in 10% sea water (Instant Ocean©, Blacksburg, Virginia, USA), and subsequently transported in aerated water to the lakes in coolers within one to three days after capture. Before release, water from the lake into which the stickleback were about to be released was gradually added to quadruple the volume of water in the cooler to avoid shocking the fish during release into the lake. They were netted out of the cooler and dropped near shore into the lake to minimize water released with the fish into the lake. The released fish showed no signs of disability, formed a single-file school and swam away from shore.

The lakes were sampled each year beginning the year after the populations were founded. Sticklebacks were captured in each lake in late May or June using unbaited Gee minnow traps mostly with 6.35 mm (1/4 inch) mesh walls. The traps were set mostly in groups of five traps at about 5 m intervals within 5 m of shore at a depth of less than 1 m near vegetation or logs and in the open. The traps were removed after about 24 hours, and all fish were anesthetized using MS222 and preserved in a 70% solution of undenatured ethanol in distilled water. The solution was replaced by fresh ethanol after about 24 hours. The preserved sticklebacks were shipped with a few drops of ethanol in plastic bags to the laboratory, where the ethanol was replaced before long-term storage in 70% ethanol and shipment to the lab.

### Library preparation and genome sequencing

All protocols for the extraction of DNA from sticklebacks and library preparation for whole genome sequencing for Scout Lake have previously been described in Kwakye et al. (2026b). We used similar protocols to extract DNA and prepare libraries for the Cheney and Warfle lake samples. Briefly, we extracted DNA using the Qiagen DNeasy 96 Blood & Tissue Kit (Germantown, Maryland) and quantified it with a Qubit Fluorometer 3.0 using Thermo Fisher Scientific High Sensitivity assay kit for dsDNA (Waltham, Massachusetts, USA). We randomly selected DNA from the Cheney sample collected in 2015 and Warfle 2024. However, we included all sticklebacks from Warfle samples collected in 2022 and 2023 since there were only 35 and 36 individuals, respectively, sampled during each of these timepoints. Library preparation was based on the plexWell™ 384 (SeqWell, Beverly, MA, USA), which is based on seqWell’s proprietary TN5 transposase and inserts Illumina i7 adapters into individual DNA samples.

### Bioinformatics Processing

We trimmed adapters with AdapterRemoval (ver. 2.2.2) (Lindgreen 2012) and mapped reads to the stickleback genome version gasAcu1-4 (Peichel et al. 2017; Roberts Kingman et al. 2021). We used Picard to add read groups and mark duplicates. We then performed base recalibration using BaseRecalibrator from GATK version 3.7 (McKenna et al. 2010; Van Der Auwera et al. 2013).

### Imputation of low-coverage data with Beagle 4.0

We imputed missing data in our low-coverage datasets using beagle 4.0 and a reference panel of 169 previously published, moderate to high coverage (9–62X coverage) genomes (Roberts Kingman et al. 2021; Kwakye et al. 2026b). The genomes in the reference panel included 69 oceanic fish, 58 putatively well-established freshwater populations, 40 from recently established populations, and two of unknown ecotype designation. We downloaded beagle 4.0 from https://faculty.washington.edu/browning/beagle/beagle.r1399.jar and performed haplotype phasing and imputation on all low-coverage data per chromosome. We ran beagle 4.0 as follows java-Xmx90G-jar beagle.27Jan18.7e1.jar gl=chrII.gz ref=WGS_170_recalibrated_Snps6mil_chrII.phased.vcf.gz map=chrII.map impute=True out=All_samples_chrII. We used GATK’s Haplotypecaller to call the genotype likelihood and used recombination maps from the ancestral Rabbit Slough population, which was previously estimated using LDhelmet (Chan et al. 2012) by (Roberts Kingman et al. 2021).

## Data Availability

All analyses were performed using custom scripts from (Kwakye 2026; Kwakye et al. 2026b). All newly generated genomes have been deposited under accession code PRJNA1472124. All previous whole genome data can be found at Sequence Read Archive (https://www.ncbi.nlm.nih.gov/sra) under accession codes PRJNA1231081[https://www.ncbi.nlm.nih.gov/sra/?term=PRJNA1231081], PRJNA671690[https://www.ncbi.nlm.nih.gov/sra/?term=PRJNA671690] and PRJNA247503[https://www.ncbi.nlm.nih.gov/sra/?term=PRJNA247503].

## Acknowledgements

R. Massengill, K. Dunker, and the Alaska Department of Fish and Game allowed us to reintroduce Threespine Stickleback to Cheney, Scout, and Warfle lakes. We also thank the Warfle and the Klein families for allowing us to introduce and sample stickleback behind their house. B. K. Barnes, C.J. Cunningham, K. L. Gould, B.P. Harris, W. Harris, B. Harris, B. Harris, A. W. Katzer, A.C. Kroska, R. Lucas, J. McMahon, E. G. Offenberg, and L. Wegner contributed assistance in the field. F. A. von Hipple provided facilities to hold sticklebacks between capture and release. This study was supported by NSF grant DEB-0919184 to MAB and NIH grant R01GM124330-01 to KRV and MAB.

