## Supplementary note for "Variable jackpot individuals provide most alleles for repeated, rapid adaptation to freshwater by anadromous Threespine Stickleback"

### Founding of Warfle lake population of Threespine Stickleback

In this paper, we compare contemporary evolution of haplotype frequencies in three populations of Threespine Stickleback (*Gasterosteus aculeatus*) that we founded in lakes using anadromous (i.e., sea-run) sticklebacks. We introduced them into Cheney Lake in 2009, Scout Lake in 2011, and "Warfle Lake" in 2019 and have sampled each lake for sticklebacks annually since the year after introduction. (See MAB's field notes, which are archived in the Ichthyology Collection, California Academy of Sciences.)

Bell et al. (2016) (Bell et al., 2016) described introduction of anadromous sticklebacks into Cheney and Scout lakes after they were treated with rotenone to exterminate Northern Pike (*Esox lucius*). This appendix describes the introduction of sticklebacks into Warfle Lake after the pike in it were exterminated.

### Choice of anadromous stickleback

This study was motivated by interest in the dynamics of genomic evolution during rapid adaptation of anadromous stickleback after they colonized freshwater. By the 1960's, it had become apparent from the geographical distribution of freshwater sticklebacks that they must have evolved from anadromous (sea-run) ancestors locally tens of thousands of times (Bell, 1976; Lindsey, 1962; Smith, 1972). However, freshwater sticklebacks exhibit a diverse set of phenotypic traits that contrast with those of their anadromous ancestors but occur in many isolated populations throughout their range. Very similar phenotypes usually must have evolved independently (i.e., convergent evolution) each time anadromous sticklebacks colonized freshwater.

The first evidence on the genetics of phenotypic convergence among geographically widespread, isolated, freshwater sticklebacks populations emerged from an elegant paper by Colosimo et al. (2005) on the developmental genetics of sticklebacks' lateral plates (Colosimo et al., 2005). Lateral plates are enlarged scales that form a single row along each side of the body, vary greatly in number, and usually differ between anadromous and freshwater sticklebacks (e.g., (Bell, 1976, 1981; Bell & Foster, 1994)). Colosimo et al. (2005) produced several results that bear on this study: (1) The *Ectodysplasin* (*EDA*) gene that strongly influences lateral plate phenotype accounts for most lateral plate morph variation (i.e. low vs. complete morph). (2) Most freshwater *EDA* alleles form a single clade relative to ancestral alleles in anadromous populations. (3) Freshwater-adaptive alleles of *EDA* occurred at low frequencies in both oceanic populations that they sampled. That has been confirmed in other populations (Bell et al., 2010), and freshwater-adaptive alleles that form separate clades at more than 300 other loci have since been identified (Jones et al., 2012; Roberts Kingman et al., 2021). (4) The freshwater-adaptive EDA alleles in almost all populations from recently deglaciated regions (i.e., <22,000 years ago) were millions of years old, much older than their freshwater habitats. That also has been confirmed for freshwater-adaptive alleles of more than 300 loci with different alleles in anadromous and freshwater sticklebacks (Jones et al., 2012; Roberts Kingman et al., 2021).

Rapid evolution of Loberg Lake sticklebacks after it was treated with rotenone and presumably recolonized naturally by anadromous sticklebacks (Bell et al., 2004) suggested that it would be possible to release anadromous sticklebacks into lakes and observe dramatic evolution within a few generations. Our experimental introductions added information on the source of the founders, the date of founding, and size of the founding population, and allowed us to observe substantial evolution using annual sampling during the first several generations in the lake. Use of anadromous sticklebacks for these three introductions simulate natural colonization of freshwater during 10 to >16 million years (Bell, 1994, 2009), Frank et al. this volume).

**Name and location of Warfle lake**

Warfle Lake is an informal name for a lake that we believe has no official name. We use the family name of the homeowners who originally allowed us to work behind their house. This lake is located south of Kasilof, Alaska on the Kenai Peninsula. The map coordinates for the release and sampling site in Warfle Lake should be used to locate the lake: 60.291° N lat., 151.3657° W long.

**Cause for absence of native sticklebacks in warfle lake.**

Warfle Lake was treated with rotenone to exterminate Northern Pike that apparently had been introduced illegally into it for sport fishing (Massengill 2022; Hendry et al. 2022). Rotenone treatment kills all fish species, and Alaska Fish and Game chose to reintroduce sticklebacks, which they agreed to delegate to one of us (MAB).

**The source population.**

We initially chose the Rabbit Slough anadromous sticklebacks population to found lake populations because it appeared to be large enough to withstand removal of 3000 adults from a run and had been characterized phenotypically (Aguirre et al., 2008). Access to Rabbit Slough is easy at the culvert under Glenn Highway and numerous sticklebacks can be caught with a few traps set in tandem at the downstream end of the culvert.

Robert Massengill, formerly of the Alaska Department of Fish and Game, permitted us to introduce sticklebacks to Warfle Lake. We used sticklebacks only from the culvert to found populations in Cheney and Scout lakes, but in 2019, we initially caught a few sticklebacks at the culvert. So we also used 30 to 50 traps to capture sticklebacks within about 40 m of the confluence of Rabbit and Palmer sloughs (Table S1), about 500 m downstream of the culvert, and at the end of South Rabbit Slough Road, to hold ancestry of the three introduced anadromous sticklebacks populations as constant as possible.

The frequency of the freshwater-adaptive allele of EDA is < 1% in oceanic populations (Bell et al., 2010; Colosimo et al., 2005), and our resources to capture and transport sticklebacks were limited. We chose to introduce ~3000 anadromous sticklebacks to each lake, including Warfle (N = 2929, Table S1), which should include about 60 freshwater-adaptive (i.e., low morph) EDA alleles. We did not know the frequencies of freshwater-adaptive alleles at other loci when we founded the first population in Cheney Lake in 2009. Subsequent estimates and evolution in our introduced populations of diverse, freshwater traits that are influenced by freshwater-adaptive alleles on multiple chromosomes, indicate that use of about 3000 sticklebacks per recipient lake captured numerous freshwater-adaptive alleles (Roberts Kingman et al., 2021).

**Capture and release.**

Anadromous (i.e., sea-run) sticklebacks were caught using nine unbaited Gee minnow traps set side by side on the bottom or in two layers of 16 traps across the downstream outlet of the culvert through which Rabbit Slough flows under Glenn Highway (61.5334°N lat., 149. 2677°W long.). Because of the weak run of anadromous sticklebacks at the culvert in June 2019, we also trapped at the juncture of Rabbit Slough and Palmer Slough (61.53429 N long., 149.26783 W lat.).

We removed live sticklebacks from the traps and placed them in coolers of aerated Rabbit Slough water. However, it can be toxic in coolers without sufficient aeration, and if the sticklebacks showed signs of hypoxia in the field, we discarded the Rabbit Slough water and replaced it with water from nearby Loberg (Junction) Lake, at the same temperature, for transport with aeration to the laboratory.

We held the Rabbit Slough sticklebacks for one to two nights in F. A. von Hippel's outdoor pool facility at the University of Alaska Anchorage. The water was deionized with enough Instant Ocean™ added to make 10% sea water and the pool water stayed below 20°C. Information on the number of fish released and dates of capture and releases are in Table S1.

After about five hours' drive from the University of Alaska Anchorage to Warfle Lake, most of the water in coolers was poured onto the ground away from the lake, and the coolers were carried down to the lake. The fish were caught with an aquarium net and released near shore. They formed a single-file school that moved rapidly away from the point of release near shore, and all of them were well oriented and swam away rapidly. A year later, their one-year old progeny were abundant near the point of release.

***Table S1:*** *Sources, numbers, and capture and release dates of anadromous Threespine Stickleback introduced into Warfle Lake. See the text for coordinates of capture and release sites. Capture and Release are dates specimens were captured in Rabbit Slough and released into Warfle Lake, respectively, in June 2019. N Culvert and N Palmer are the number of* *stickleback caught in Rabit Slough at the culvert and at the juncture of Rabbit and Palmer sloughs on each date, and N Total is the number of stickleback from both sites each date. Totals for each capture site and their sum are shown on the bottom line.*

| Capture | N Culvert | Palmer N | Total | Total |
|---|---|---|---|---|
| 6 | 81 | 0 | 81 | 8 |
| 7 | 112 | 321 | 433 | 8 |
| 8 | 143 | 285 | 428 | 9 |
| 12 | 311 | 272 | 583 | 15 |
| 14 | 286 | 404 | 690 | 15 |
| 16 | 300 | 35 | 335 | 18 |
| 17 | 145 | 234 | 379 | 18 |
| Total | 1378 | 1551 | 2929 | |

**ACKNOWLEDGEMENTS**

We thank F. A. von Hippel for use of his outdoor pools to hold sticklebacks before release. E. and J. Warfle allowed us to release sticklebacks and sample sticklebacks for several years before J. and D. Ames acquired their house and permitted us to continue sampling. A. P. Hendry released some of the sticklebacks to Warfle Lake. B. K. Barnes, K. L. Gould, several undergraduates from Stony Brook University and the University of Alaska Anchorage, and members of D. M. Kingsley's laboratory, who are too numerous to list, helped found and make annual samples from Cheney and Warfle Lakes. Supported by NSF grant DEB-0919184 to MAB and NIH grant 1R01GM124330-01 to KRV and MAB.
